# A new insight into underlying disease mechanism through semi-parametric latent differential network model

**DOI:** 10.1101/397265

**Authors:** Yong He, Jiadong Ji, Lei Xie, Xinsheng Zhang, Fuzhong Xue

## Abstract

**Background:** In genomic studies, to investigate how the structure of a genetic network differs between two experiment conditions is a very interesting but challenging problem, especially in high-dimensional setting. Existing literatures mostly focus on differential network modelling for continuous data. However, in real application, we may encounter discrete data or mixed data, which urges us to propose a unified differential network modelling for various data types.

**Results:** We propose a unified latent Gaussian copula differential network model which provides deeper understanding of the unknown mechanism than that among the observed variables. Adaptive rank-based estimation approaches are proposed with the assumption that the true differential network is sparse. The adaptive estimation approaches do not require precision matrices to be sparse, and thus can allow the individual networks to contain hub nodes. Theoretical analysis shows that the proposed methods achieve the same parametric convergence rate for both the difference of the precision matrices estimation and differential structure recovery, which means that the extra modeling flexibility comes at almost no cost of statistical efficiency. Besides theoretical analysis, thorough numerical simulations are conducted to compare the empirical performance of the proposed methods with some other state-of-the-art methods. The result shows that the proposed methods work quite well for various data types. The proposed method is then applied on gene expression data associated with lung cancer to illustrate its empirical usefulness.

**Conclusions:** The proposed latent variable differential network models allows for various data-types and thus are more flexible, which also provide deeper understanding of the unknown mechanism than that among the observed variables. Theoretical analysis, numerical simulation and real application all demonstrate the great advantages of the latent differential network modelling and thus are highly recommended.

## Background

In genomic studies, graphical model has been an important tool to capture dependence among different genes. Particularly, Gaussian graphical model has been widely applied to infer the relationship between genes at the transcriptional level [1–4]. Under the Gaussian assumption, estimating the structure of the graphical model is equivalent to recover the support of precision matrix which is defined to be the inverse of the covariance matrix. However, in some cases, compared to focusing on a particular network, it is of greater interest to investigate how the network of connected gene pairs change from one experimental condition to another, which provides deeper insights on an underlying biological process such as identification of pathways that correspond to such a change. For instance, medical experiment usually involves two groups: the patient group and the control group. The analysis of group difference in biological networks or pathways may offer us a new insight into the underlying disease mechanism, which have extensive biomedical and clinical applications, such as identifying effective targets for drug development in a cost-effective and timely manner. Indeed, differential networking modelling has recently emerged as an important tool to analyze a set of changes in graph structure between two conditions [see, for example 5–17]. In the context of genomic analysis, it is reasonable to assume that two genes are defined to be connected in the differential network if the magnitude of their conditional dependency relationship changes between two conditions. The precision matrix which is defined as the inverse of covariance matrix can capture the conditional dependency relationship. Thus the differential network is typically modelled as the difference of two precision matrices and this type of modelling has been widely used [7– 9, 14, 15]. Figure 1 (a),(b),(c) illustrate the definition of differential network. Each node represents a gene. For two groups depicted in (a) and (b), there is an edge between genes (*i, j*) if and only if (*i, j*)-th element of **Ω** is nonzero. For each edge, there exists a weight which is the magnitude of (*i, j*)-th element of **Ω**. Gene pair (*i, j*) is defined to be connected in the differential network in (c) if the magnitudes of (*i, j*)-th elements of two precision matrices change between two groups.

**Figure 1.**
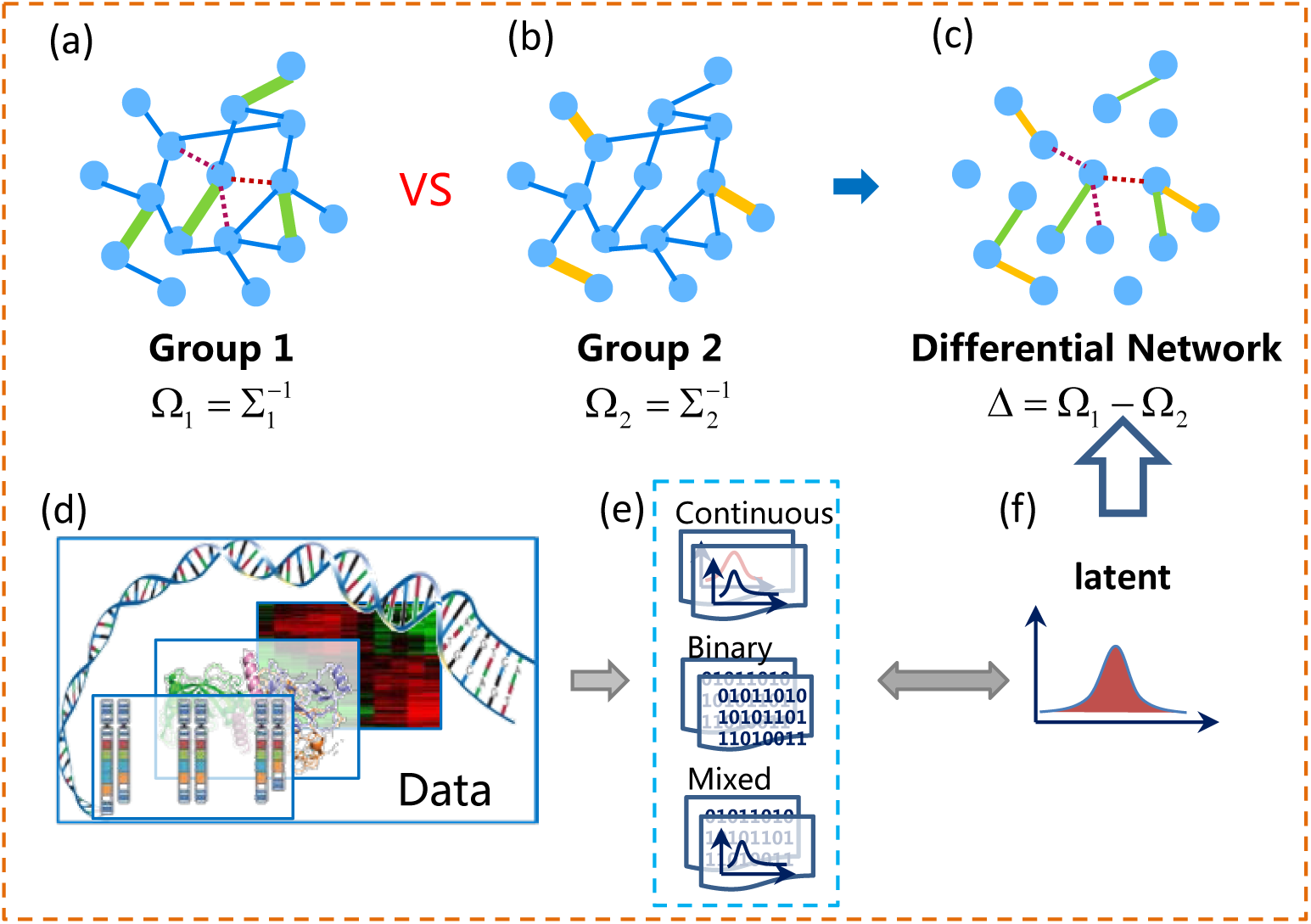
Illustration of latent differential network.

One straightforward approach to estimate the difference of two precision matrices is to separately estimate the precision matrices and then subtract the estimates. In the high dimensional setting where the dimension *p* is much larger than the sample size *n*, which is often the case for genomic study, many estimation approaches for the precision matrix have been proposed and proved to enjoy nice theoretical properties and computation advantage under the key assumption of sparsity. And this topic has been an active area of research in recent years [18–22].

Another type of approach to estimate the difference of two precision matrices is to jointly estimate the precision matrices. Guo et al. [23] penalized the joint loglikelihood with a hierarchical penalty that targets the removal of common zeros in the inverse covariance matrices across categories. Danaher et al. [24] proposed the joint graphical Lasso, which is based upon maximizing a penalized log-likelihood with generalized fused Lasso or group Lasso penalty. Motivated by the constrained *𝓁*_1_ minimization approach to precision matrix estimation of [22], Zhao et al. [7] proposed an estimation approach to directly estimate the difference of the precision matrices.

For the separately estimating methods, Liu et al. [25] proposed the nonparanormal family to relax the Gaussian assumption. While the nonparanormal family is much larger than the standard parametric Gaussian family, the independence relations among the variables are still encoded in the precision matrix. In addition, Liu et al. [26] proposed a semiparametric approach called nonparanormal SKEPTIC to estimate high dimensional undirected graphical models efficiently and robustly and proved that the nonparanormal SKEPTIC achieves the optimal parametric rates of convergency in terms of precision matrix estimation and graph recovery. Xue and Zou [27] proposed a similar regularized rank-based estimation idea for estimating nonparanormal graphical models and analyzed adaptive versions of rank-based Dantzig selector and clime estimators. He et al. [28] proposed a multiple testing procedure to estimate high-dimensional nonparanormal graphical model and proved that the proposed procedure can control the false discovery rate (FDR) asymptotically.

The disadvantage of Gaussian or nonparanormal graphical models lies in that they are only tailored for modeling continuous data. However, in genomic studies, we may encounter discrete data (e.g. CNV data and SNP data), continuous data (e.g. gene expression and methylation data) or data of hybrid types with both discrete and continuous variables. Besides, in some circumstances, even if the data are continuous, we still need to transform the data into discrete data to remove the heterogeneity (e.g. batch effect, outliers and population stratification). For instance, in the analysis of gene expression data collected from different platforms, to remove the unwanted variation among different experiments known as the batch effects, numerical expression data are often transformed into 0/1 binary data, where lower expression values are encoded as 0 and higher expression values are encoded as 1. In this setting, it is reasonable to assume that the discrete variable is obtained by discretizing a latent variable. Fan et al. [29] proposed a general model named the latent Gaussian copula graphical model, assuming that the observed discrete data are generated by discretizing a latent continuous variable at some unknown cutoff.

In this paper, we consider estimating differential network for various types of biological data in a joint way. We propose a unified semi-parametric latent variable differential network model. The latent differential network model is illustrated in Figure 1 (e)-(f). For biological data, there exist continuous data, discrete data or data of hybrid types with both continuous and discrete data. It is assumed that these data are collected by transforming latent continuous variables which are unobservable. We are interested in the differential network of the latent variables, which provide deeper understanding of the unknown mechanism than that among the observed variables. To the best of our knowledge, our work provides the first method for differential network estimation for binary or mixed data with theoretical guarantees under the high dimensional scaling. The advantages of the proposed methods lie in the following aspects: (I) Our method provides a way to infer the differential network structure among latent variables, which provides deeper understanding of the unknown mechanism than that among the observed variables. (II) Theoretical analysis shows that the proposed methods achieve the same parametric rates of convergence for both difference matrix estimation and differential graph recovery, as if the latent variables were observed. (III) The proposed methods are much more robust to outliers due to the rank-based correlation matrix estimator. (IV) The proposed approaches do not require precision matrices to be sparse, and thus can allow the individual networks to contain hub nodes. Simulation result shows that the proposed method performs much better and more robustly than several state-of-the-art methods. The proposed methods are applied on a gene expression data set associated with lung cancer. A target gene WIF1 stands out by the proposed method, which indeed is verified as a frequent target for epigenetic silencing in various human cancers [30]. The real data example illustrates the great usefulness of the current work.

## Methods

In this part, we propose novel definitions of latent differential network model for various types of data. In essence, we define the differential network as the difference of two precision matrices of the latent variables, which greatly generalizes the applicability in areas such as bioinformatics, medical research and so on.

### Gaussian copula differential graphical model

We first review the definition of the Gaussian copula distribution. Let *f* = {*f*_1_, …, *f*_*p*_} be a set of strictly increasing univariate functions. A *p* dimensional random variable ***X*** = (*X*_1_, …, *X*_*p*_)^*T*^ is said to follow the Gaussian copula distribution if and only if *f* (***X***) := (*f*1 (*X*1), …, *p* (*X p*))^*T*^ := ***Z*** *∼ N p* (***µ***, **Σ**) and is noted as ***X*** *∼* NPN(***µ***, **Σ**, *f*), where ***µ*** = (*µ*_1_, …, *µ*_*p*_), **Σ** = [Σ_*jk*_] are respectively the mean vector and the correlation matrix of the Gaussian variate ***Z***. The conditional independence structure of ***X*** is encoded by the sparsity pattern of **Ω** = **Σ**^*-*1^. Specifically, it can be shown that *X*_*i*_ is conditionally independent of *X*_*j*_ given all other variables if and only if *ω*_*ij*_ = 0, where *ω*_*ij*_ is the (*i, j*) th element of **Ω**. Therefore, the differential network of the Gaussian copula variables can be defined to be the difference between the two precision matrices, just the same as for the parametric Gaussian case.

Assume ***X***_*i*_ = (*X*_*i*1_, …, *X*_*ip*_)^*T*^ for *i* = 1, …, *n*_*X*_ are independent observations of the expression levels of *p* genes from one group denoted by ***X*** and ***Y***_*i*_ = (*Y*_*i*1_, …, *Y*_*ip*_)^*T*^ for *i* = 1, …, *n*_*Y*_ from the other denoted by ***Y***, ***X*** *∼*NPN(***µ***^*X*^, **Σ**^*X*^, *f* ^*X*^) and ***Y*** *∼*NPN(***µ***^*Y*^, **Σ**^*Y*^, *f* ^*Y*^). The differential network is defined to be the difference between the two precision matrices, denoted by **Δ**_0_ = **Ω**^*Y*^ *-***Ω**^*X*^, where **Ω**^*Y*^ and **Ω**^*X*^ are the inverse matrices of **Σ**^*Y*^ and **Σ**^*X*^ separately.

We propose a rank-based estimator of **Σ**^*X*^. It is known that if ***Z*** ∼ NPN(***µ***, **Σ**, *f*), then we have.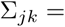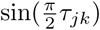, where *τ*_*jk*_ is Kendall’s tau correlation between *Z*_*j*_ and *Z*_*k*_. Thus we can estimate the unknown correlation matrix **Σ**^*X*^ by:

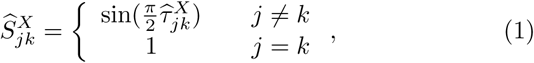

Where 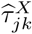 is the sample Kendall’s tau correlation between *X*_*j*_ and *X*_*k*_. Similarly, we can estimate **Σ**^*Y*^ in the same way and obtain the estimator ***Ŝ***^*Y*^. Motivated by the direct estimation method of the difference of two precision matrices proposed by [7], one can obtain the estimator of **Δ**_0_ by solving

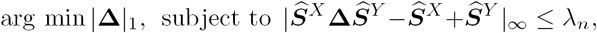

which is equivalent to the optimization problem:

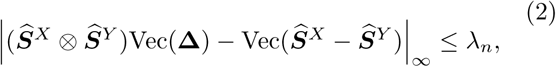

where ⊗ denotes the Kronecker product, *|***Δ***|*_1_ = **Σ** _*jk*_*δ*_*jk*_ is the element-wise *𝓁*_1_ norm of the matrix **Δ**. Here, for a matrix ***A*** = [*A*_*jk*_], *|****A****|*_*∞*_ = max_*jk*_ *|A*_*jk*_*|* and for a vector ***a*** = (*a*_*j*_), *|****a****|*_*∞*_ = max_*j*_ *|a*_*j*_*|*.

As seen from Equation (2), the proposed approach can directly estimate the difference matrix without implicitly estimating the individual precision matrices. Thus there is no need to assume the sparsity of (**Σ**^*Y*^)^*-*1^ and (**Σ**^*X*^)^*-*1^. We only need to assume that **Δ**_0_ is sparse. Besides, compared to the sample covariance matrix, the rank-based estimators here can enjoy modelling flexibility and estimation robustness, especially when outliers exist.

### Latent Gaussian copula differential graphical model for binary data

In the analysis of gene expression data, to remove the batch effects, numerical expression data are often transformed into 0/1 binary data, where lower expression values are encoded as 0 and higher expression values are encoded as 1. To estimate the underlying differential network for the binary data from two different groups, we assume that the observed discrete data are generated by discretizing a latent continuous variable at some unknown cutoff. To make the model more flexible, we assume the latent continuous variable is Gaussian copula distributed instead of Gaussian. Let ***B*** = (*B*_1_, *B*_2_, *…, B*_*p*_)^*T*^ *∈* {0, 1}^*p*^ be a *p*-dimensional 0*/*1-random vector. The 0*/*1-random vector ***B*** satisfies the latent Gaussian copula model (LGCM) for binary data, if there exists a *p* dimensional random vector ***X*** *∼* NPN(**0**, **Σ**, *f*) such that

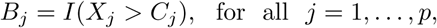

where *I*(*·*) is the indicator function and the cutoff ***C*** =(*C*_1_, *…, C*_*p*_) is a vector of constants. Then we denote ***B*** *∼* LGCM(**Σ**, *f,* ***C***). We call **Σ** the latent correlation matrix. The latent Gaussian copula model involves parameters (**Σ**, *f,* ***C***). Merely based on the binary random vector ***B***, only *f*_*j*_(*C*_*j*_), *j* = 1, *…, p* are identifiable. Denote **Λ** = (Λ_1_, …, Λ_*p*_), where Λ_*j*_ = *f*_*j*_(*C*_*j*_). For notational simplicity, we write LGCM(**Σ**, **Λ**) for LGCM(**Σ**, *f,* ***C***).

Assume 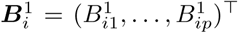 for *i* = 1, *…, n*_1_ are independent observations of the binary expression levels of *p* genes from one group denoted by ***B***^1^ and 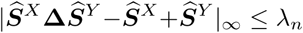 for *i* = 1, *…, n*_2_ from the other denoted by ***B***^2^, where ***B***^1^ *∼*LGCM(**Σ**^1^, **Λ**^1^) and ***B***^2^ *∼*LGCM(**Σ**^2^, **Λ**^2^). The differential network is defined to be the difference between the two precision matrices, denoted by 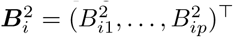 Motivated by Equation (2), we should first derive estimators for **Σ**^1^ and **Σ**^2^. For ease of presentation, we only present the procedure to construct the estimator for **Σ**^1^, estimator for **Σ**^2^ can be obtained similarly. Denote the Kendall’s tau correlation between 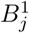 and 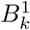 by 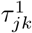, it can be shown that 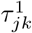 satisfies:

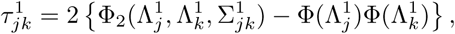

Where

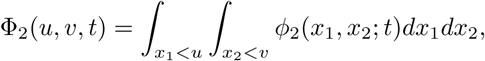

is the cumulative distribution function of the standard bivariate normal distribution, *ϕ*_2_(*x*_1_, *x*_2_; *t*) is the probability density function of the standard bivariate normal distribution with correlation *t*. Denote by

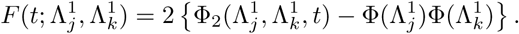

For any fixed 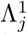 and 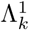, it can be shown that 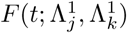 is a strictly monotonic increasing function on *t ∈* (*-*1, 1) and thus is invertible. Given 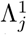 and 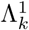, one can estimate 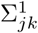 by 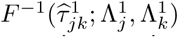. However, the cutoff values are unknown in practice. As 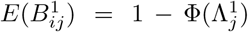, we can estimate 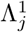 by 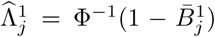, where 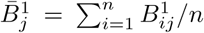 Thus the Kendall’s tau rank-based correlation matrix estimator 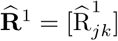 for **Σ**^1^ is a *p × p* matrix with element entry given by

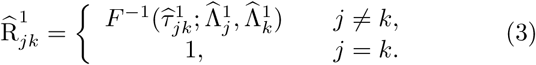

Similarly, the Kendall’s tau rank-based correlation matrix estimator 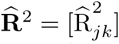 for **Σ**^2^ is a *p × p* matrix with element entry given by

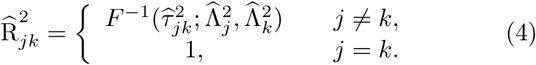

Motivated by Equation (2), we can obtain an estimator of 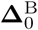 by solving the following optimization problem:

arg min |Δ|_1_; subject to

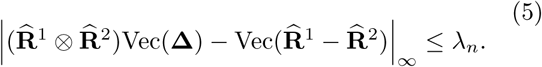

For the binary data, we aim to infer the differential network among latent variables, which provides deeper understanding of the unknown mechanism than that among the observed binary variables. Thus, our model complements the existing work on high dimensional differential network estimation, which mostly focused on learning differential network among observed variables including, for example, the Ising model.

### Latent Gaussian copula differential graphical model for Mixed data

In the analysis of biological data, there also exists the case where some biological data are discrete while some others are continuous. For instance, multi-level omics data integrative analysis involves gene mutation, expression, methylation, metabolome and phenome data. In this case, mixed data appear naturally. We start with the definition of the latent Gaussian copula model for mixed data. Assume that ***M*** = (***M***_1_, ***M***_2_), where ***M***_1_ represents the *p*_1_-dimensional binary variables and ***M***_2_ represents the *p*_2_-dimensional continuous variables. The random vector ***M*** satisfies the latent Gaussian copula model for mixed data, if there exists a *p*_1_ dimensional random vector ***X***_1_ such that ***X*** = (***X***_1_, ***M***_2_) *∼* NPN(0, **Σ**, *f*) and

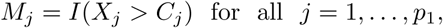

where ***C*** = (*C*_*1*_, *…, C* _*P*1_) is a vector of constants. Then we denote ***M*** *∼* LGCM(0, **Σ**, *f,* ***C***), and call **Σ** the latent correlation matrix. In the latent Gaussian copula regression model, the binary components ***M***1 are generated by a latent continuous random vector ***X***_1_ truncated at ***C***, and combining with the continuous components ***M***_2_, ***X*** = (***X***_1_, ***M***_2_) satisfies the Gaussian copula model. For the binary data ***M***_1_, onlyΛ_*j*_ = *f*_*j*_(*C*_*j*_), *j* = 1, *…, p*_1_ are identifiable. For the continuous components ***M***_2_, the marginal transformations *f*_*j*_(*·*), *j* = *p*_1_ + 1, *…, p* are identifiable.

Assume 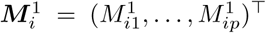 for *i* = 1, *…, n*_1_ are independent observations of the expression levels of *p* genes from one group denoted by ***M*** ^1^ and 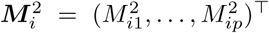 for *i* = 1, *…, n*_2_ from the other denoted by ***M*** ^2^, where ***M*** ^1^ *∼* LGCM(**Σ**^1^, **Λ**^1^) and ***M*** ^2^ *∼* LGCM(**Σ**^2^, **Λ**^2^). The differential network is defined to be the difference between the two precision matrices, denoted by 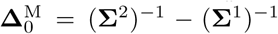 Similar to the discussions in the last sections, we first need to construct estimators for **Σ**^1^ and **Σ**^2^. For ease of presentation, we only present the procedure to construct the estimator for **Σ**^1^, estimator for **Σ**^2^ can be obtained similarly. For discrete components 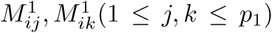, as what we have discussed in the last subsection with a slight change of notation, we can estimate 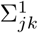 by:

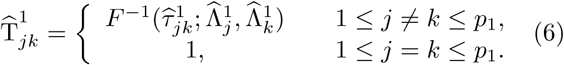

For continuous components 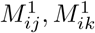, as what we have discussed, we can estimate 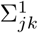 by:

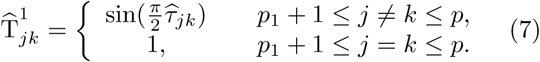

where 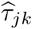is defined as follows:

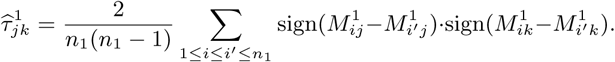

We still need to consider the mixed case. Without loss of generality, we assume that 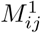 is binary and 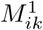 is continuous. In this case, the Kendall’s tau correlation can be expressed by

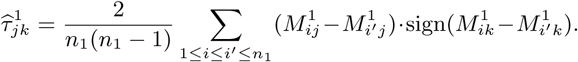

The population version of Kendall’s tau correlation 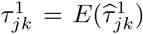 can be expressed by 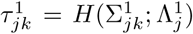, where

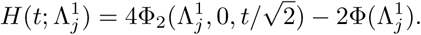

Moreover, for fixed 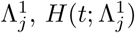, is an invertible function of *t*. The parameter 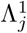 could be estimated by 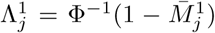, where 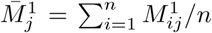. Thus when 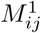 is binary and 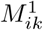 is continuous, 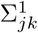 can be estimated by the Kendall’ tau rank-based estimator:

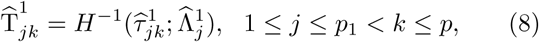

Where 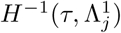 is the inverse function of 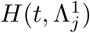 for fixed 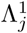. Thus the Kendall’s tau rank-based correlation matrix estimator 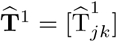 for **Σ**^1^ is a *p×p* matrix with corresponding element entry given by Equation (6), (7), and (8) respectively. Similarly, we can obtain estimator 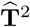 for **Σ**^2^. Motivated by Equation (2), we can obtain an estimator of **Δ**_0_ by solving the following optimization problem:

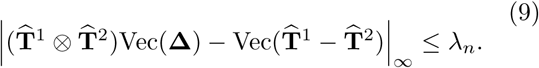

We show that the rank-based covariance matrix estimators achieve the same parametric rate of convergence for both difference matrix estimation and differential graph recovery in the Additional file 1. Thus the extra modelling flexibility comes at almost no cost of statistical efficiency. Besides, for the binary data or data of hybrid types with both binary and continuous variables, the differential network among latent variables can be well estimated, which provides deeper understanding of the unknown mechanism than that among the observed variables.

### Implementation

In this section we will present how to solve the optimization problems in Equation (2), (5), and (9). For ease of presentation, we only present the procedure to obtain the solution to optimization problem in Equation (2) and solutions to optimization problems in Equation (5) and Equation (9) can be obtained in the similar way.

Recall that in Equation (2), the optimization problem is

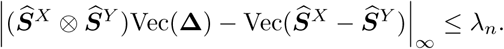

Let **Δ** = [*δ*_*jk*_]_1*≤j,k≤p*_ and define ***θ*** to be the *p*(*p* + 1)*/*2 × 1 vector with ***θ*** = (*δ*_*jk*_)_1*≤j≤k≤p*_. Estimating a symmetric **Δ** is thus equivalent to estimating ***θ***, which alleviates the computation burden especially when *p* is large. Define the *p*^2^ *×p*(*p*+1)*/*2 matrix **Γ** with columns indexed by 1 *≤ j ≤ k ≤ p* and with rows indexed by *l* = 1, *…, p* and *m* = 1, *…, p*, so that each entry is labeled by Γ_*lm,jk*_. For *j ≤ k*, let Γ_*jk,jk*_ = Γ_*kj,jk*_ = 1 and set all other entries of **Γ** equal to zero. With these notations, one may consider the following optimization problem:

*θ̂* = arg min |*θ*|_1_ subject to

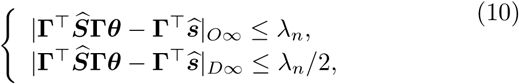

where ***Ŝ*** = ***Ŝ***^*X*^ *⊗* ***Ŝ***^*Y*^, ***ŝ*** = Vec(***Ŝ***^*X*^ *-* ***Ŝ***^*Y*^) and for a *p*(*p* + 1)*/*2 *×* 1 vector ***c***, *|****c****|*_*O∞*_ denotes the sup-norm of the entries of ***c*** corresponding to the off diagonal elements of its matrix form, while *|****c****|*_*D∞*_ denotes the sup-norm of the entries of ***c*** corresponding to the diagonal elements. The matrix form of 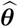 will be denoted by 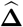 in the following sections. The optimization problem in Equation (10) can be solved by the alternating direction method of multipliers (ADMM), for a thorough discussion, we refer to [31]. For the optimization problem in Equation (10), to apply the ADMM algorithm, we rewrite it as:

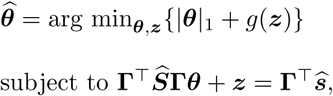

where the function *g*(*·*) is defined by

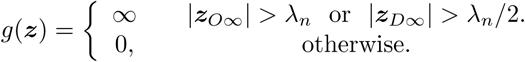

The augmented Lagrangian can be written as

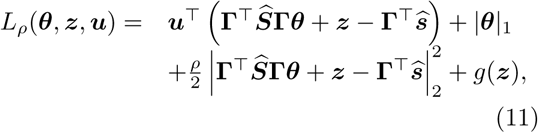

where ***u*** is the Lagrange multiplier and *ρ* is a positive penalty parameter which can be specified by users. The ADMM algorithm is based on minimizing the augmented Lagrangian in (11) over ***θ*** and ***z*** and then applying a dual variable update to the Lagrange multiplier ***u***, which yields the updates

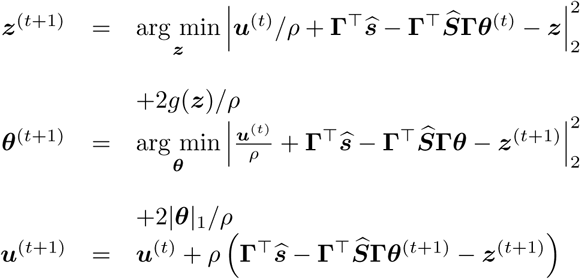

for iterations *t* = 0, 1, 2 *…*. As for the tuning parameter *λ*_*n*_ in (10), it can be chosen by an approximate Akaike information criterion (AIC). *λ*_*n*_ is chosen to minimize

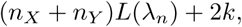

where *k* is the effective degrees of freedom that can be approximated by 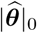 and *L*(*λ*_*n*_) represents the loss function either *L*_*∞*_ or *L*_F_ which are defined by

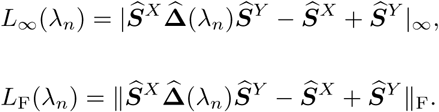

In this paper we focus on the loss functions with the supremum and Frobenius norms for further theoretical development. One may also use other matrix norms, such as spectral norm:

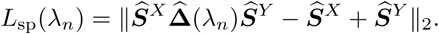

Similarly, for the latent Gaussian copula model for binary data, one can solve the following optimization problem:

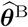 = arg min |*θ*|_1_ subject to

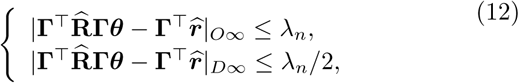

Where 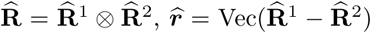. The matrix form of 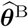 will be denoted by 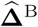 in the following sections. For the latent Gaussian copula model for mixed data, one can solve the following optimization problem:

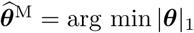 subject to

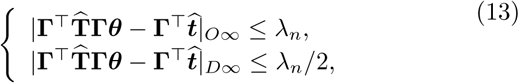

Where 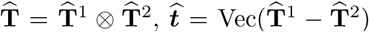. The matrix form of 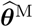 will be denoted by 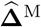 in the following sections. Besides, corresponding Akaike information criterion can be proposed to choose the tuning parameter *λ*_*n*_.

### Simulation

**Simulation for Gaussian copula differential graphical model** In this part, we conduct simulation study for differential network estimation under Gaussian copula model. We mainly focus on the graphs that contain hub nodes. First we generate the edge set E^*X*^ for the group ***X***. We partition *p* features into 5 equally-sized and non-overlapping sets: *C*_1_ *∪ C*_2_ *· · · ∪ C*_5_ = {1, *…, p*}, *|C*_*k*_*|* = *p/*5, *C*_*i*_ *∩ C*_*j*_ = *∅*. For the smallest *i ∈C*_*k*_, we set (*i, j*) *∈ C*_*k*_ for all *{j i* : *j ∈ C*_*k*_. *}* The non-zero entries of **Ω**^*X*^ is then determined by the edge set E^*X*^, where **Ω**^*X*^ = (**Σ**^*X*^)^*-*1^. Next, the value of each nonzero entry of **Ω**^*X*^ was generated from a uniform distribution with support [*-*0.75, *-*0.25] *∪* [0.25, 0.75]. To ensure positive definiteness of **Ω**^*X*^, let **Ω**^*X*^ = **Ω**^*X*^ + (0.2 + *|λ*_min_(**Ω**^*X*^) *|*)**I**. At last the **Ω**^*X*^ is rescaled such that **Σ**^*X*^ is a correlation matrix. Then we proceed to generate the differential network **Δ**_0_. We randomly select two hub nodes from the 5 equally-sized and non-overlapping sets. The differential network **Δ**_0_ is generated such that the connections of these two hub nodes change sign between **Ω**^*X*^ and **Ω**^*Y*^. The correlation matrix **Σ**^*X*^ and **Σ**^*Y*^ are generated by (**Ω**^*X*^)^*-*1^ and (**Ω**^*Y*^)^*-*1^ respectively. Finally we generate *n*_*X*_ i.i.d observations of ***Z***^*X*^ from the *N* (**0**, **Σ**^*X*^) distribution and *n*_*Y*_ i.i.d observations of ***Z***^*Y*^ from the *N* (**0**, **Σ**^*Y*^) distribution. Next we sample *n*_*X*_ i.i.d samples from the nonparanormal distribution NPN(**0**, **Σ**^*X*^, *f* ^*X*^) and *n*_*Y*_ i.i.d samples from the nonparanormal distribution NPN(**0**, **Σ**^*Y*^, *f* ^*Y*^). For simplicity, we use the same univariate transformations on each dimension: 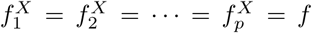 and *f* ^*X*^= *f* ^*y*^sample data from the nonparanormal distribution, we also need *g* := *f* ^*-*1^. We consider the Gaussian CDF Transformation of *g* which is used in [26].

In the simulation study, we let *p* = 50,80,100,120 and *n*_*X*_ = *n*_*Y*_ = 100. The simulation result is based on 100 replications. For each simulated data set, we apply three estimation methods. That is, the direct differential network estimator (DDN) in [7], the rank-based differential network estimator (RDN) and the direct differential network estimator based on the latent variable ***Z*** and Pearson correlation (ZP-DDN). In ZP-DDN, we assume that ***Z***^*X*^ and ***Z***^*Y*^ are observed and the Pearson correlation estimator of cov(***Z***^*X*^) and cov(***Z***^*Y*^) are plugged into the direct estimation procedure. While ZP-DDN are often not available in real applications, we use ZP-DDN as benchmarks for quantifying the information loss of the remaining estimators.

We evaluate the performance of the estimation methods from two aspects: support recovery and estimation error. The support recovery results are evaluated by true positive rate (TPR) and true negative rate (TNR) along a range of tuning parameter *λ*. Suppose the true difference matrix **Δ**_0_ has the support 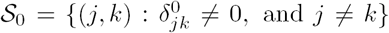 and its estimator 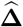 has the support set **ŝ**. TPR and TNR are defined as follows:

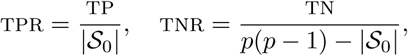

where TP and TN are the numbers of true positives and true negatives respectively, which are defined as

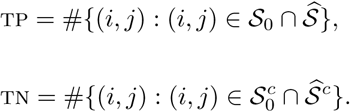

To evaluate the support recovery performance, we use the true discovery rate, which is defined as TD = TP*/| Ŝ* _0_*|*. As for the estimation error, we calculate the element-wise *L*_*∞*_ norm and Frobenius norm of 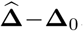

**Simulation for latent Gaussian copula differential graphical model** In this part, we conduct simulation study for differential network estimation under Latent Gaussian copula model. We assume that the cutoff vector ***C*** *∼* Unif [0, 1] and let **Σ**^1^ and **Σ**^2^ be generated in the same way as **Σ**^*X*^ and **Σ**^*Y*^ described in the last subsection. We consider the following three Scenarios:

- **Scenario 1** Generate data 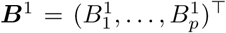, where 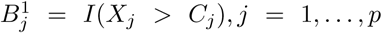 and ***X*** *∼* NPN(**0**, **Σ**^1^, *f* ^1^); Generate data 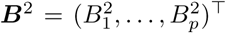, where 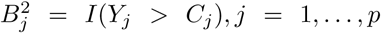 and ***Y*** *∼* NPN(**0**, **Σ**^2^, *f* ^2^). The transformation functions *f* ^1^ and *f* ^2^ are Gaussian CDF transformation.
- **Scenario 2** Generate data 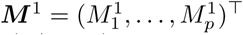, where 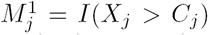, *j* = 1,…*p* ***X*** *∼* NPN(0, **Σ**^1^, *f* ^1^) and 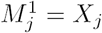, *j* = 1,…,*p* Generate data 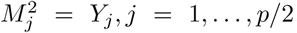, where 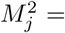 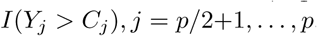, and ***Y*** *∼* NPN(0, **Σ**^2^, *f* ^2^) and 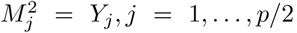. The transformation functions *f* ^1^ and *f* ^2^ are Gaussian CDF transformation.
- **Scenario 3** Generate data 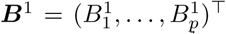, where 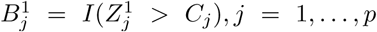 and ***Z***^1^ *∼ N* (**0**, **Σ**^1^), where 10 entries in each ***Z***^1^ is randomly sampled and replaced by −5 or 5; Generate data 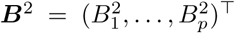, where 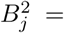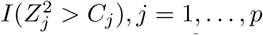 and ***Z***^2^ *∼ N* (**0**, **Σ**^2^), where 10 entries in each ***Z***^2^ is randomly sampled and replaced by −5 or 5.

In Scenario 1 and Scenario 3, we generate binary data. Scenario 1 corresponds to the latent Gaussian copula model and Scenario 3 corresponds to the setting where the binary data can be misclassified due to the outliers of the latent Gaussian variable. Scenario 3 is designed to investigate the robustness of the proposed approach. Scenario 2 corresponds to the mixed data generated from the latent Gaussian copula model.

### Application to gene expression data sets related to lung cancer

In this section we consider the differential network estimation for a gene expression data set related to lung cancer. The data set is publicly available from the Gene Expression Omnibus at accession number GDS2771 and was studied in [24]. It includes 22,283 microarray-derived gene expression measurements from large airway epithelial cells sampled from 97 patients with lung cancer and 90 controls in the data set. It is of interest to investigate how the structure of the gene co-expression network differs between the group of patients with lung cancer and the control group. It may shed light on underlying lung cancer mechanisms. In this real example study, we limited our analysis to the 122 genes in the Wnt signaling pathway. The Wnt signaling pathway has recently emerged as a critical pathway in lung carcinogenesis as already demonstrated in many cancers and particularly in colorectal cancer [32]. The Gene expression levels were analyzed on a logarithmic scale. Each gene feature was standardized to have mean zero and standard deviation 1 within the cancer samples and the controls separately.

## Results

### Simulation results for Gaussian copula differential graphical model

The receiver operating characteristic (ROC) curves of the three estimation methods are depicted in Figure 2. It shows that the proposed method RDN compares favourably with the benchmark method ZPDDN, which means that the information loss is negligible. Besides, Figure 2 also shows that DDN performs pretty bad in the non-Gaussian case.

**Figure 2.**
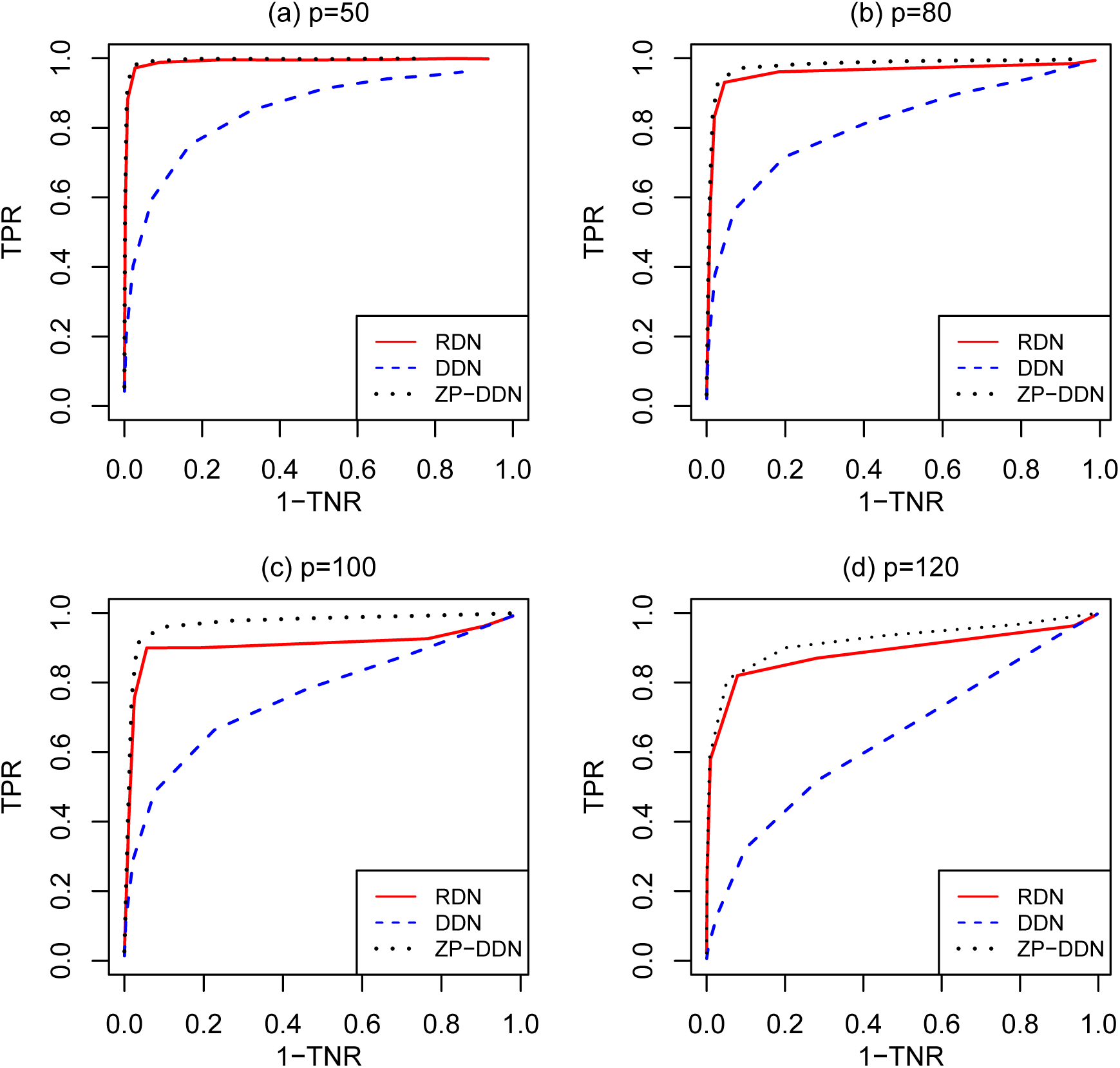
Receiver operating characteristic curves under Gaussian copula model with dimensionality varying from 50 to 120. The red line represents the proposed RDN method, the black dotted represents the benchmark method ZP-DDN, the blue dotted line represents DDN method.

Table 1 gives the true discovery rates with different loss functions. The results also show the method RDN compares favourably with the benchmark method ZPDDN. For all the methods, tuning using the *L*_F_ gives better true discovery rates than tuning using the *L*_*∞*_. Table 1 depicts the elementwise *L*_*∞*_ norm estimation accuracies of the thresholded estimators tuned using the loss functions *L*_*∞*_ and *L*_F_. From Table 1, we can see that the *L*_F_ loss function gives slightly better results than the *L*_*∞*_ loss function. For all the methods, the elementwise *L*_*∞*_ norm estimation accuracy are comparable. We point out that it is possible for RDN to simultaneously give better support recovery but similar estimation than DDN. The reason is that estimation error depends on the magnitudes of the estimated entries, while support recovery depends only on whether the entries are nonzero. Besides, RDN has comparable performance with the benchmark method ZP-DDN in terms of both support recovery and estimation accuracy, which indicates that the information loss of the estimator RDN is negligible.

**Table 1.** Average true discovery rates (%) and average estimation errors over 100 simulations.

| Average true discovery rates |  |  |  |  |  |  |
| --- | --- | --- | --- | --- | --- | --- |
| $p$ | ZP-DDN | | RDN | | DDN | |
| | $L_\infty$ | $L_F$ | $L_\infty$ | $L_F$ | $L_\infty$ | $L_F$ |
| 50 | 74.0 (13.6) | 83.2 (10.9) | 75.6 (14.0) | 89.1 (11.3) | 45.9 (24.7) | 27.8 (17.3) |
| 80 | 91.4 (16.4) | 99.6 (4.3) | 95.2 (14.2) | 100.0 (0.0) | 44.9 (34.6) | 51.0 (42.8) |
| 100 | 96.3 (14.1) | 100.0(0.0) | 99.5 (5.2) | 100.0 (0.0) | 39.3 (40.3) | 50.0 (49.1) |
| 120 | 78.8 (16.8) | 100.0(0.0) | 79.0 (18.2) | 100.0 (0.0) | 23.4 (41.3) | 30.0 (46.3) |
| Average estimation errors in the elementwise $L_\infty$ norm | | | | | | |
| $p$ | $L_\infty$ | $L_F$ | $L_\infty$ | $L_F$ | $L_\infty$ | $L_F$ |
| 50 | 3.26 (0.41) | 2.91 (0.33) | 3.08 (0.32) | 2.59 (0.35) | 2.27 (0.12) | 2.41 (0.21) |
| 80 | 2.06 (0.28) | 1.92 (0.06) | 1.98 (0.21) | 1.91 (0.00) | 1.97 (0.09) | 1.94 (0.08) |
| 100 | 1.86 (0.15) | 1.82 (0.00) | 1.82 (0.04) | 1.82 (0.00) | 1.87 (0.10) | 1.83 (0.04) |
| 120 | 1.12 (0.17) | 0.87 (0.00) | 1.12 (0.18) | 0.87 (0.00) | 0.89 (0.07) | 0.87 (0.00) |

### Simulation results for Latent Gaussian copula differential graphical model

The ROC curves for Scenario 1 and Scenario 2 with different dimensionality *p* (varying from 50 to 120) is presented in Figure 3. Table 2 give the true discovery rates with different loss functions and the elementwise *L*_*∞*_ norm estimation accuracies of the thresholded estimators tuned using the loss functions *L*_*∞*_ and *L*_F_, respectively. For method ZR-RDN, we assume that the latent Gaussian copula variables are observed. In particular, the rank-based correlation matrix estimator of the latent Gaussian copula variables are plugged into the direct estimation procedure. With a slight abuse of notation, the RDN method here refers to either the rank-based method for binary data or for mixed data. The ROC curves in Figure 3 show that the rank-based methods RDN proposed for latent Gaussian copula model (binary and mixed) perform pretty well even when the dimensionality is larger than the sample size.

By the ROC curves in Figure 4, we can find that RDN is more robust to the data misclassification than the benchmark estimator ZP-DDN. The robustness of RDN to outliers illustrates the advantage of the dichotomization method. In the absence of misclassification, it is seen that the ROC curves of RDN and ZR-RDN are similar, which indicates little information loss for differential network recovery due to the dichotomization procedure. Table 3 gives the true discovery rates with different different loss functions for Scenario 3 and presents the elementwise *L*_*∞*_ norm estimation accuracies of the thresholded estimators tuned using the loss functions *L*_*∞*_ and *L*_F_ for Scenario 3. From Table 3, we can see that the *L*_F_ loss function gives slightly better results than the *L*_*∞*_ loss function. Besides, we can see that the elementwise *L*_*∞*_ norm estimation accuracy are comparable. This is also true for Scenario 1 and Scenario 2.

**Table 2.** Simulation results over 100 replications for Scenario 1 and Scenario 2.

| Average true discovery rates(%) |  |  |  |  |
| --- | --- | --- | --- | --- |
| $p$ | Scenario 1 | | Scenario 2 | |
| | $L_\infty$ | $L_F$ | $L_\infty$ | $L_F$ |
| 50 | 78.8 (15.2) | 98.4 (5.9) | 79.6 (13.8) | 40.8 (25.6) |
| 80 | 76.4 (23.1) | 100.0(0.0) | 83.4 (17.0) | 88.2 (17.6) |
| 100 | 89.5 (22.1) | 100.0(0.0) | 84.8 (20.0) | 99.3 (3.9) |
| 120 | 76.5 (31.0) | 94.0(24.0) | 82.4 (15.2) | 100.0 (0.0) |
| Average estimation errors in the elementwise $L_\infty$ norm | | | | |
| $p$ | Scenario 1 | | Scenario 2 | |
| | $L_\infty$ | $L_F$ | $L_\infty$ | $L_F$ |
| 50 | 2.66 (0.26) | 2.21 (0.15) | 3.23 (0.40) | 3.85 (0.55) |
| 80 | 2.10 (0.20) | 1.91 (0.00) | 2.29 (0.35) | 2.14 (0.32) |
| 100 | 1.88 (0.13) | 1.82 (0.00) | 2.03 (0.28) | 1.83 (0.08) |
| 120 | 1.00 (0.16) | 0.87 (0.00) | 1.17 (0.16) | 0.88 (0.07) |

**Table 3.** Simulation results over 100 replications for Scenario 3.

| $p$ | Average true discovery rates(%) | | | | | |
| --- | --- | --- | --- | --- | --- | --- |
|  | ZP-DDN |  | RDN |  | ZR-RDN |  |
| | $L_{\infty}$ | $L_F$ | $L_{\infty}$ | $L_F$ | $L_{\infty}$ | $L_F$ |
| 50 | 39.8 (39.5) | 46.1 (47.3) | 87.6 (14.7) | 97.3 (7.4) | 88.0 (11.0) | 90.0 (12.4) |
| 80 | 32.4 (41.9) | 35.5 (47.9) | 80.7 (14.8) | 99.8 (2.5) | 89.5 (8.7) | 95.4 (7.2) |
| 100 | 23.5 (40.2) | 31.7 (46.9) | 75.6 (20.3) | 100.0(0.0) | 84.0 (12.0) | 99.1 (4.2) |
| 120 | 16.0 (37.0) | 16.0 (37.0) | 52.9 (44.6) | 68.0(47.1) | 70.4 (26.8) | 93.0 (24.8) |
| Average estimation errors in the elementwise $L_{\infty}$ norm | | | | | | |
| $p$ | $L_{\infty}$ | $L_F$ | $L_{\infty}$ | $L_F$ | $L_{\infty}$ | $L_F$ |
| 50 | 2.15 (0.03) | 2.16 (0.01) | 2.05 (0.15) | 2.12 (0.08) | 2.05 (0.17) | 2.02 (0.15) |
| 80 | 1.91 (0.02) | 1.91 (0.01) | 1.91 (0.12) | 1.92 (0.04) | 1.92 (0.12) | 1.91 (0.08) |
| 100 | 1.82 (0.02) | 1.82 (0.00) | 1.88 (0.12) | 1.82 (0.00) | 1.90 (0.12) | 1.83 (0.04) |
| 120 | 0.87 (0.00) | 0.87 (0.00) | 0.91 (0.09) | 0.87 (0.00) | 0.97 (0.11) | 0.88 (0.05) |

**Figure 3.**
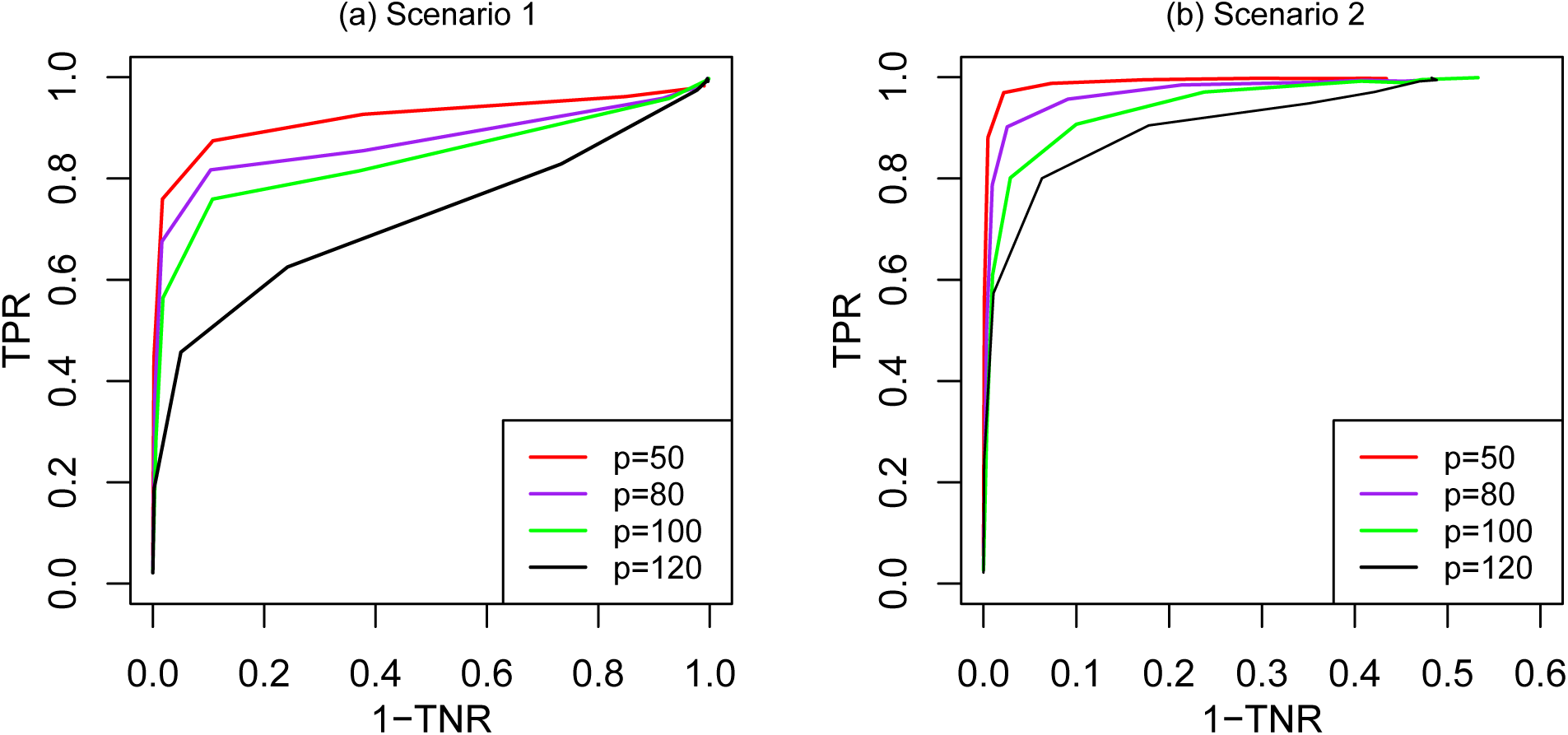
Receiver operating characteristic curves for Scenario 1 and Scenario 2 under latent Gaussian copula model, with dimensionality varying from 50 to 120.

**Figure 4.**
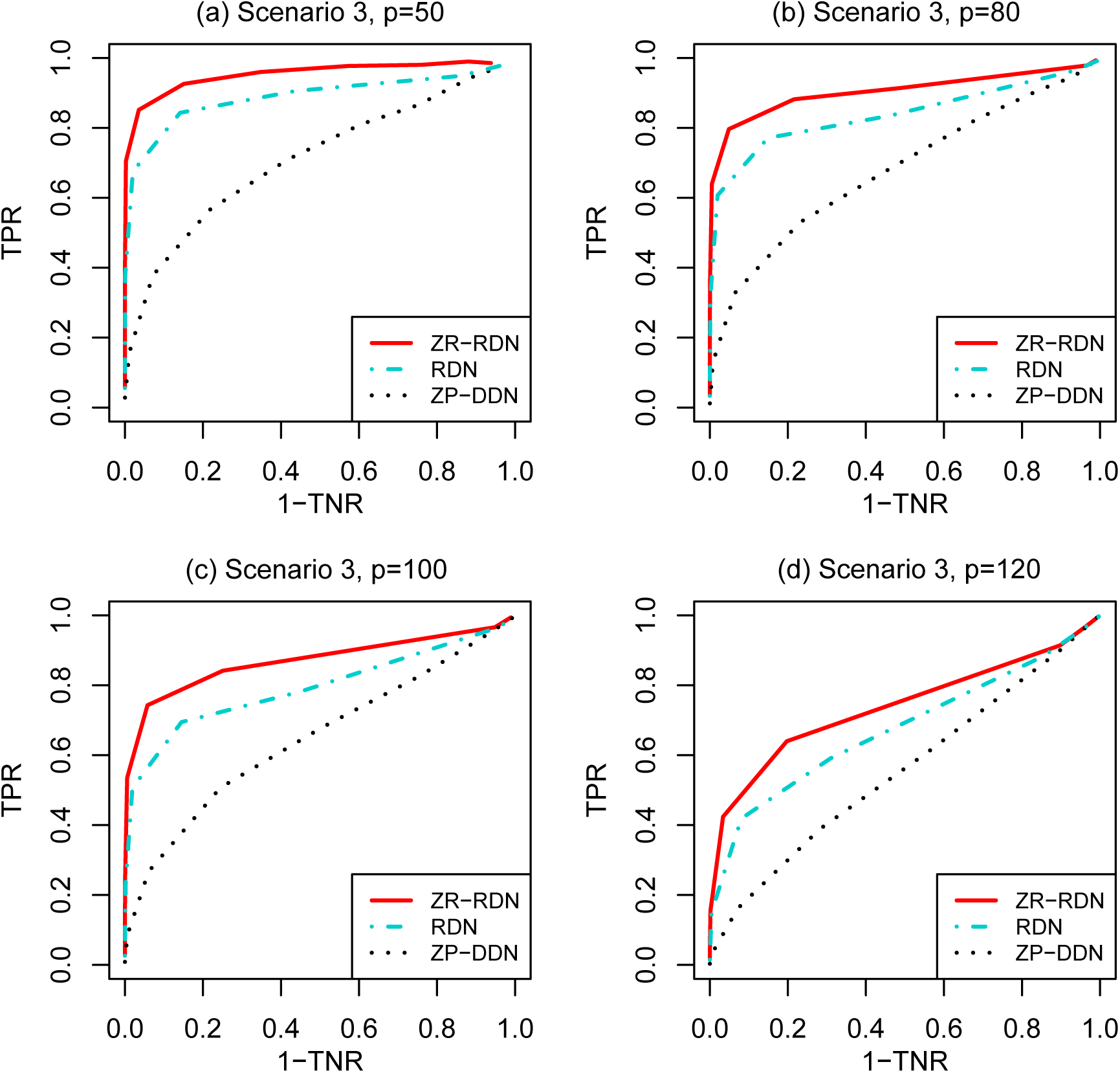
Receiver operating characteristic curves for Scenario 3 under latent Gaussian copula model, with dimensionality varying from 50 to 120. The red line represents the proposed RDN method, the black dotted represents the benchmark method ZP-DDN, the blue dotted line represents DDN method.

### Theoretical results

The estimators 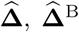 and 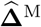, after an additional threshold step, are shown to be able to recover not only the support of the true **Δ**_0_ but also the signs of its nonzero entries as long as those entries are sufficiently large. Besides, under mild conditions, the estimation errors bounds in terms of matrix Frobenius norm and elementwise *𝓁*_*∞*_ norm both achieve the parametric rate 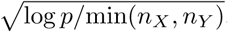, see details in Additional file 1. It indicates that the extra modeling flexibility and robustness come at almost no cost of statistical efficiency and it seems as if the latent variable can be observed. Thus these new estimators can be used as a safe replacement of Gaussian estimators even when the data are truly Gaussian. Compared to the separate and joint approaches to estimating differential networks [e.g. 22, 23] which require sparsity on each **Σ**^*-*1^, the proposed direction estimation methods for different types of data only require the sparsity of the difference matrix **Δ**_0_. The detailed theorems and proofs are in the Additional file 1 available online.

### Results of application

In the real application part, we compare three estimation methods. The first method is the Gaussian copula RDN method, which we denote as C-RDN. The second method is the latent Gaussian copula RDN method, which we denote as B-RDN. In specific, we first apply the adaptive dichotomization method implemented by the ArrayBin package in R to remove the batch effect in the gene expression data. The adaptive dichotomization method transforms the numerical gene expression data into 0/1 binary data. The genes with high expression level are encoded as 1 and the genes with lower expression level are encoded as 0. Then we apply the B-RDN to the 0/1 binary data. The third method is the direct differential network estimation method proposed by [7] with Gaussian assumption, which we denote as DDN.

We conduct Shapiro-Wilk test on the gene data set and 63% of the genes reject the normality null hypothesis. Therefore, the Gaussian assumption of DDN method is violated in this real data example. Thus we expect that C-RDN which relaxes the Gaussian assumption may provide a more reliable result. The deficiency of the C-RDN method lies in that it does not take the batch effect of the genes expression data from different platforms into consideration. For the B-RDN method, it removes the batch effect.

Figure 5 depicts the differential network estimated by the three methods. Table 4 gives the hub genes selected out by different estimation methods. For method C-RDN, the tuning parameter *λ* is selected by the AIC criterion with the elementwise *𝓁*_1_ norm loss function. To ensure a fair comparison, the tuning parameter *λ* for method B-RDN and DDN are selected such that the number of edges in the estimated differential graphs by all three methods are almost the same. The number of edges selected by the three methods are 56, 59 and 52, respectively. From Figure 5, we can see that B-RDN identifies an obvious hub gene WIF1 that is an extracellular antagonist of WNT. WIF1 is a frequent target for epigenetic silencing in various human cancers [30]. WIF1 promoter is frequently methylated in non-small cell lung cancer (NSCLC) cells to down-regulate its mRNA expression [33]. Both C-RDN and B-RDN select out a common hub gene APC. APC expression in lung cancer are associated with survival time and is also related to cancer metastasis [34]. Both C-RDN and DDN select out a common hub gene, MAPK8, which plays a significant role in the promotion of lung inflammation and tumorigenesis subsequent to tobacco smoke exposure [35]. The expression level of DVL2 was reported significantly higher in lung adenocarcinomas than in squamous carcinomas, and was associated with poor tumor differentiation [36]. Winn et al. [37] reported that the restoration of FZD9 signaling inhibited both cell proliferation and anchorage-independent growth, promoted cellular differentiation, and reversed the transformed phenotype in NSCLC. The overexpression of MMP7 was associated with tumor proliferation, and a poor prognosis in NSCLC [38]. RAC1 generally plays an important role in cancer progression and metastasis [39].

**Table 4.** Hub genes selected by different methods.

| DDN | PRKACA | MAPK8 | CACYBP | CAMK2B | SFRP1 | CSNK2A2 | TCF7 |
| --- | --- | --- | --- | --- | --- | --- | --- |
|  | BTRC | RUVBL1 |  |  |  |  |  |
| C-RDN | PLCB2 | DVL2 | MAPK8 | PLCB1 | APC | WNT2 | FZD9 |
|  | WNT11 | DKK1 | SFRP4 |  |  |  |  |
| B-RDN | WIF1 | MMP7 | RAC1 | LEF1 | APC | PRKACA | WNT8B |
|  | BAMBI |  |  |  |  |  |  |

**Figure 5.**
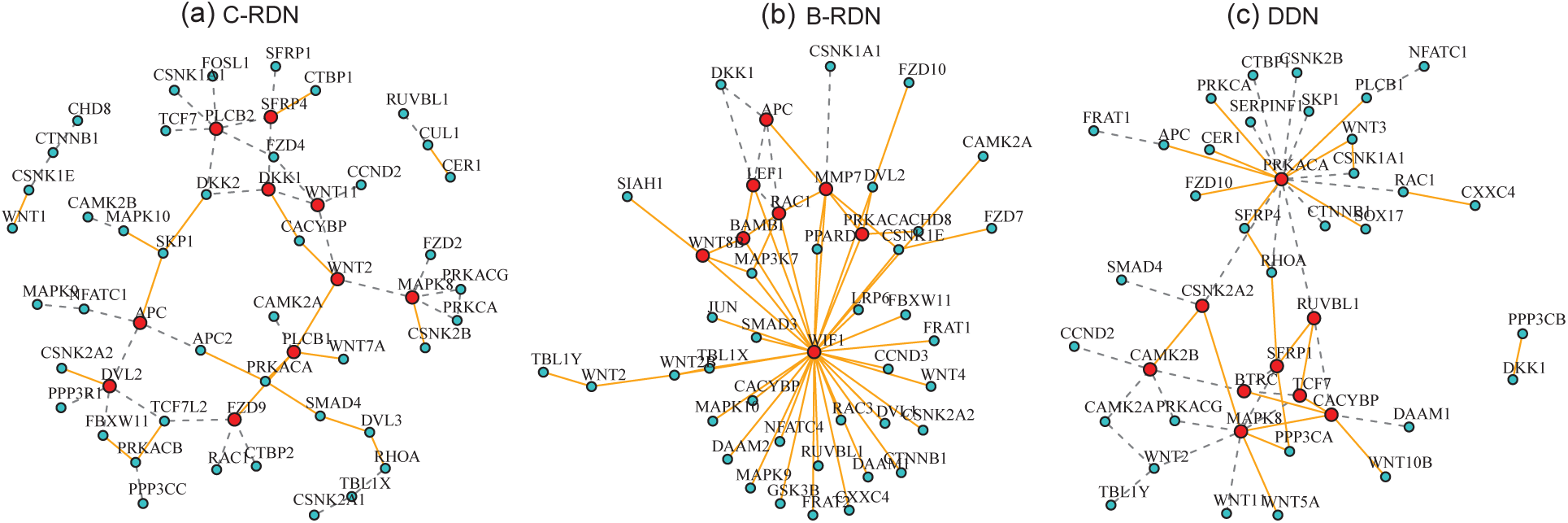
Differential network estimated by different methods. Orange edges show an increase in conditional dependency from control group to lung cancer patient group; grey edges show a decrease. Red points stand for hub genes which have edges with more than 3 other genes.

By comparing (a) and (b) in Figure 5, we can see that the estimated differential network can be very different with/without considering the batch effect. Although it is inevitable to result in information loss in the discretization procedure for method B-RDN, [40] argued that this procedure can potentially improve the accuracy of the statistical analysis. In real data example, we recommend to use the B-RDN method to remove the batch effect despite the little information loss. At last we argue that statistical comparison of group difference in this biological network or pathway can provide new insight into the underlying lung cancer mechanism, which may further offer more effective targets for drug development.

To further interpret the underlying biological implications of the identified hub genes, we conducted Gene Ontology (GO) enrichment analysis. Table 5 shows the common GO terms enriched by C-RDN, B-RDN and DDN. The GO enrichment analysis is performed using R package “clusterProfiler” with the P-value adjusted by Benjamini-Hochberg method. It shows that our methods (C-RDN, B-RDN) have smaller P-value than DDN. The common molecular function and cellular component suggest that the change of frizzled binding, Wnt-protein binding and beta-catenin destruction complex are important in the etiology of lung cancer. These predictions are supported by the literatures [41– 43], which indicates that the proposed differential network model can provide biological meaningful underlying signals.

**Table 5.**
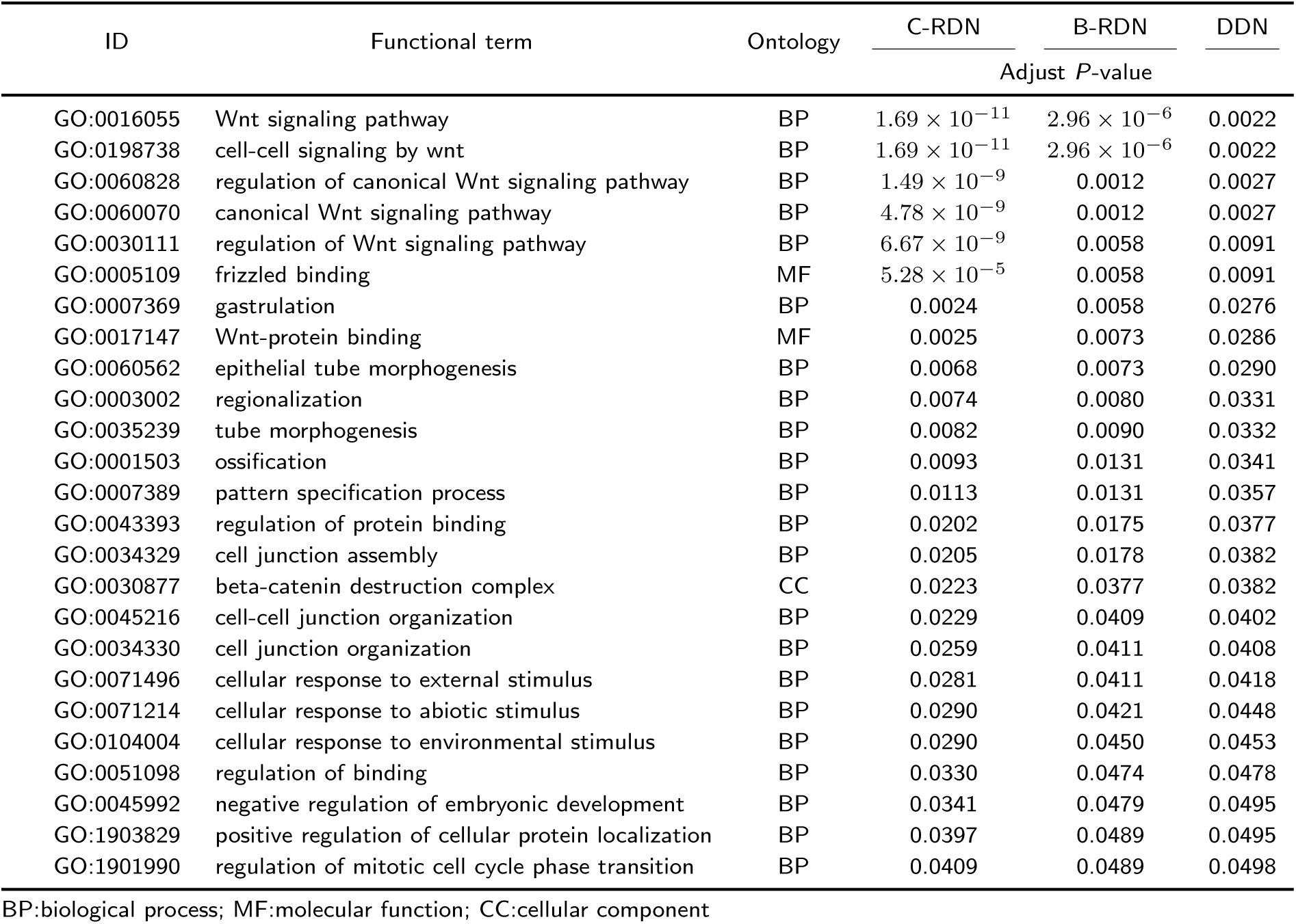
Gene Ontology (GO) enrichment analysis result BP:biological process; MF:molecular function; CC:cellular component

## Discussion and conclusions

A complex disease phenotype (e.g. diabetes, cancer) often reflects various pathobiological processes that interact in a network rather than the abnormality of a single gene. Such interactions are not static processes, instead they are dynamic in response to changing genetic, epigenetic and environmental factors, which further entails the analysis of differential network. In this paper, we propose adaptive estimation approaches for latent variable differential network model with the assumption that the true differential network is sparse, which do not require precision matrices to be sparse. The latent variable differential network model is fundamentally different from the existing ones in the literature in the sense that the differential structure in the unobserved latent variables are of primary interest. Theoretical analysis shows that the proposed methods achieve the same parametric convergence rate for both the difference of the precision matrices estimation and differential structure recovery, which means that the extra modelling flexibility comes at almost no cost of statistical efficiency. The unified latent variable differential network model provides deeper understanding of the unknown genomic mechanism than that among the observed variables.

The current work could be extended in the following two aspects. First, in this paper, we consider the following optimization problem to directly estimate the difference matrix **Δ**:

arg min |Δ|_1_, subject to

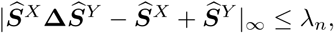

where ***Ŝ*** ^*X*^ and ***Ŝ*** ^*Y*^ denote the rank-based estimators of the covariance matrices. The D-trace loss function [15, 44] can also be applied to to directly estimate the precision matrix difference. Thus, we may also consider the D-trace loss function to estimate the Gaussian copula and latent Gaussian copula differential graphical models. In specific, the difference matrix **Δ** could be eatimated by:

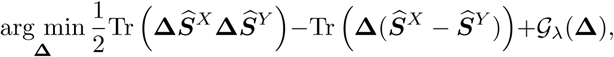

where *λ >* 0 is a regularization parameter and *𝒢* _*λ*_ is a decomposable non-convex penalty function which has the form *𝒢* _*λ*_ = **Σ** _*j,k*_ *g*_*λ*_(Δ_*jk*_), such as smoothly clipped absolute deviation (SCAD) penalty [45]. The theoretical guarantees are still needed to be investigated, but we expect that the empirical performance could be comparable.

Second, for the latent Gaussian copula differential graphical model, we focus on the binary data. In fact, the methods can be extended to the discrete data with more than two categories. The properties of this procedure are left for future investigation as there are a lot of work still needed to be done.

The proposed latent variable differential network models are very flexible and provide deeper understanding of the unknown biological mechanism. It is demonstrated latent differential network models enjoy great advantages over existing models and thus are highly recommended in real application.

## Additional File

### Additional file 1 —Supplementary Files

Contains the theoretical guarantee of of the proposed methods and proofs. (PDF 284 kb)

### Author’s contributions

YH and JDJ contributed to the study design, analytical preparation and the writing of the manuscript. YH and JDJ performed the simulation studies. JDJ analyzed the data, XL, XSZ and FZX revised the manuscript. All authors read and approved this version of the manuscript.

### Availability of data

The gene expression data set related to lung cancer is publicly available from the Gene Expression Omnibus at accession number GDS2771.

## Acknowledgements

This work was supported by grants from the Natural Science Foundation of Shandong Province (ZR2018BH033) and National Natural Science Foundation of China (grant number 81803336, 11571080 and 81573259). The funding body played no role in the design, writing or decision to publish this manuscript.

## Abbreviations

SNP:Single Nucleotide Polymorphisms; CNV: Copy Number Variation; ROC: Receiver Operating Characteristic; TPR: True Positive Rate; TNR: True Negative Rate; TDR: True Discovery Rate; FDR: False Discovery Rate; GO: Gene Ontology.

## Ethics approval and consent to participate

Not applicable.

## Consent for publication

Not applicable.

